# Synthetic hypermutation: gene-drug mutation rate synergy reveals a translesion synthesis mechanism

**DOI:** 10.1101/086512

**Authors:** Romulo Segovia, Yaoqing Shen, Scott A. Lujan, Steven Jones, Peter C. Stirling

## Abstract

Gene-gene or gene-drug interactions are typically quantified using fitness as readout because the data is continuous and easily measured in high-throughput. However, to what extent fitness captures the range of other phenotypes that show synergistic effects is usually unknown. Using *Saccharomyces cerevisiae*, and focusing on a matrix of DNA repair mutants and genotoxic drugs, we quantify 76 gene-drug interactions based on both mutation rate and fitness and find that these parameters are not necessarily overlapping. Independent of fitness defects we identified six cases of synthetic hypermutation, where the combined effect of the drug and mutant on mutation rate was greater than predicted. One example occurred when yeast lacking *RAD1* were exposed to cisplatin and we characterized this interaction using whole-genome sequencing. Our sequencing results indicate mutagenesis by cisplatin in *rad1*Δ cells appeared to depend almost entirely on interstrand crosslinks at GpCpN motifs. Interestingly, our data suggests that the 3’ base in these motifs templates the addition of the mutated base. This result differs from cisplatin mutation signatures in XPF-deficient *C. elegans* and supports a model in which translesion synthesis polymerases perform a slippage and realignment extension across from the damaged base. Accordingly, DNA polymerase ζ activity was essential for mutagenesis in cisplatin treated *rad1*Δ cells. Together these data reveal the potential to gain new mechanistic insights from non-fitness measures of gene-drug interactions and extend the use of mutation accumulation and whole-genome sequencing analysis to define DNA repair mechanisms.

## SIGNIFICANCE

Cancer cells often have defects in DNA repair and are killed effectively by drugs that damage DNA. However surviving cells can acquire additional mutations after treatment with these genotoxic chemicals. Here we apply a simple model system to reveal synergy between specific DNA repair mutations and genotoxic drugs that occurs independently of fitness defects. Moreover, by analyzing the entire genome of a mutagenized cell population we identify a signature of mutations that informs the mechanism of the translesion synthesis DNA damage tolerance pathway. Our work establishes a conceptual framework for predicting the mutational burden of cells surviving genotoxin treatment and demonstrates the utility of model organism mutation signature analysis for generating mechanistic insights.

## INTRODUCTION

Genome maintenance pathways suppress the accumulation of mutations derived from chemical lesions or mismatches in DNA that arise during normal metabolic processes. Despite thousands of potentially mutagenic lesions occurring per cell per day, mitotic cell division exhibits extremely low rates of mutation (10^-8^ – 10^-10^per bp per generation) under normal conditions (1). Cells with defects in DNA repair pathways are generally more permissive for mutation accumulation (MA) and this likely underlies the predisposition of individuals inheriting cellular DNA repair defects to cancer formation (2). Increasing the rate of mutations in a cell population makes it more likely that the necessary mutations in oncogenes and tumour-suppressor genes will arise in a given lineage leading to cellular transformation and proliferation (3, 4).

While DNA repair defects can predispose to cancer formation, when acquired somatically they may serve as an Achilles Heel that can be exploited for cancer therapy. This is because chemotherapeutic chemicals often work by damaging DNA, stalling DNA replication or disrupting mitosis, all of which could be potentially deleterious to a cell with an acquired DNA repair deficiency (2). Several well-documented examples of this type of gene-drug interaction exist for the BRCA1 and 2 genes where their mutational status can predict sensitivity of tumors to cisplatin and its derivatives (5). Indeed there are now multiple efforts underway to inhibit additional DNA repair proteins themselves to further sensitize cancers to killing with genotoxic agents or to overwhelm tumor cells with already debilitated DNA repair capacity(2).

In cancer, as in model systems, there are typically surviving cells after a genotoxic insult. These cells bear a signature of mutations associated with the genotoxin they survived (6, 7). Mutation signature analysis of tumor genomes has been refined in the past 5 years and a set of 30 canonical mutation signatures is now maintained in the COSMIC database (7, 8). In some cases the etiology of mutation patterns is strongly linked to specific genetic mutations or environmental exposures, while in other cases the etiology remains unknown. Studies in model organisms have sought to dissect which aspects of a mutation signature are due to specific deficiencies in genome maintenance factors or to specific chemical treatments (9-11). Indeed, the largest such study to date in *C. elegans* characterized both a panel of mutant strains and the effects of Aflatoxin B1, mechlorethamine and cisplatin (11).

The intention of genotoxin treatments clinically is to kill cells rather than mutagenize them. Model organism studies have also provided a means to map genetic networks underlying genotoxin sensitivity. The systematic identification of synthetic lethal interactions or chemical-genetic interactions has been led by studies in budding yeast, *Saccharomyces cerevisiae*. Indeed, a full pairwise gene-gene interaction study is now complete for both essential and non-essential yeast genes (12). In addition several thousand small molecules have been profiled for sensitivity and resistance across the yeast knockout (YKO) collections (13). These approaches are being combined to understand the effects of chemical perturbations on genetic interaction networks and identifying gene-gene synergies in drug sensitivity (14, 15). In each of these studies the primary readout for synergy between genes and chemicals is fitness as it is quantitative, simple to measure in high-throughput and informative. Nevertheless, other quantitative phenotypic readouts are possible and the YKO collection has been profiled by numerous biochemical, cytological and functional phenotypes (16).

Reasoning that DNA repair deficiencies would result in cell death, mutagenesis of survivors, or both after a genotoxic insult we assessed the overlap of fitness and mutagenesis for representative chemical genotoxins in yeast cells defective for all major DNA repair pathways. Quantifying growth and mutation rates showed little overlap between these parameters and further revealed cases of unexpected hypermutation. We characterize one such interaction between *RAD1* and cisplatin by whole-genome sequencing and uncover evidence of a novel translesion synthesis mechanism for yeast DNA polymerase ζ (Rev3). Together these data define gene-drug interactions in a new way, underscore a novel mutation signature in yeast and apply experimental MA and whole genome sequencing to suggest new DNA repair mechanisms.

## RESULTS

### A network of genotoxin-induced fitness defects and mutation rates

To investigate gene-drug interactions, we first established a panel of DNA repair mutants representing the major DNA repair pathways in yeast **(Table 1)**. Haploid yeast bearing gene deletions impairing homologous recombination (HR), non-homologous end joining (NHEJ), nucleotide-excision repair (NER), base-excision repair (BER), mismatch repair (MMR), translesion synthesis (TLS), Fanconi Anemia-like (FA) and post-replication repair (PRR) pathways were chosen to represent loss or deficiencies of each pathway. To account for the multiple steps of, and routes for, DSB repair by HR we also included exo1Δ, *rad52*Δ, *sgs1*∆, *mus81*∆ and *mph1*∆ lines as these mutants specifically impair end resection, strand exchange, double Holliday junction dissolution, resolution or other HR steps respectively(17). This panel of wildtype (WT) and 12 mutant strains **(Table 1)** was exposed to five classes of DNA damaging agents, represented by the bifunctional alkylator cisplatin, the antimetabolite fluorouracil (5FU), the topoisomerase I inhibitor camptothecin (Cpt), the topoisomerase II inhibitor etoposide (Etp), and the monofunctional alkylator methyl methanesulfonate (MMS). We screened 78 pairwise gene-drug combinations plus WT for changes in fitness and, in 76 cases, were also able to measure mutation rates (*rad5*Δ growth was impaired to a degree that prevented fluctuation analysis).

**Table 1.**
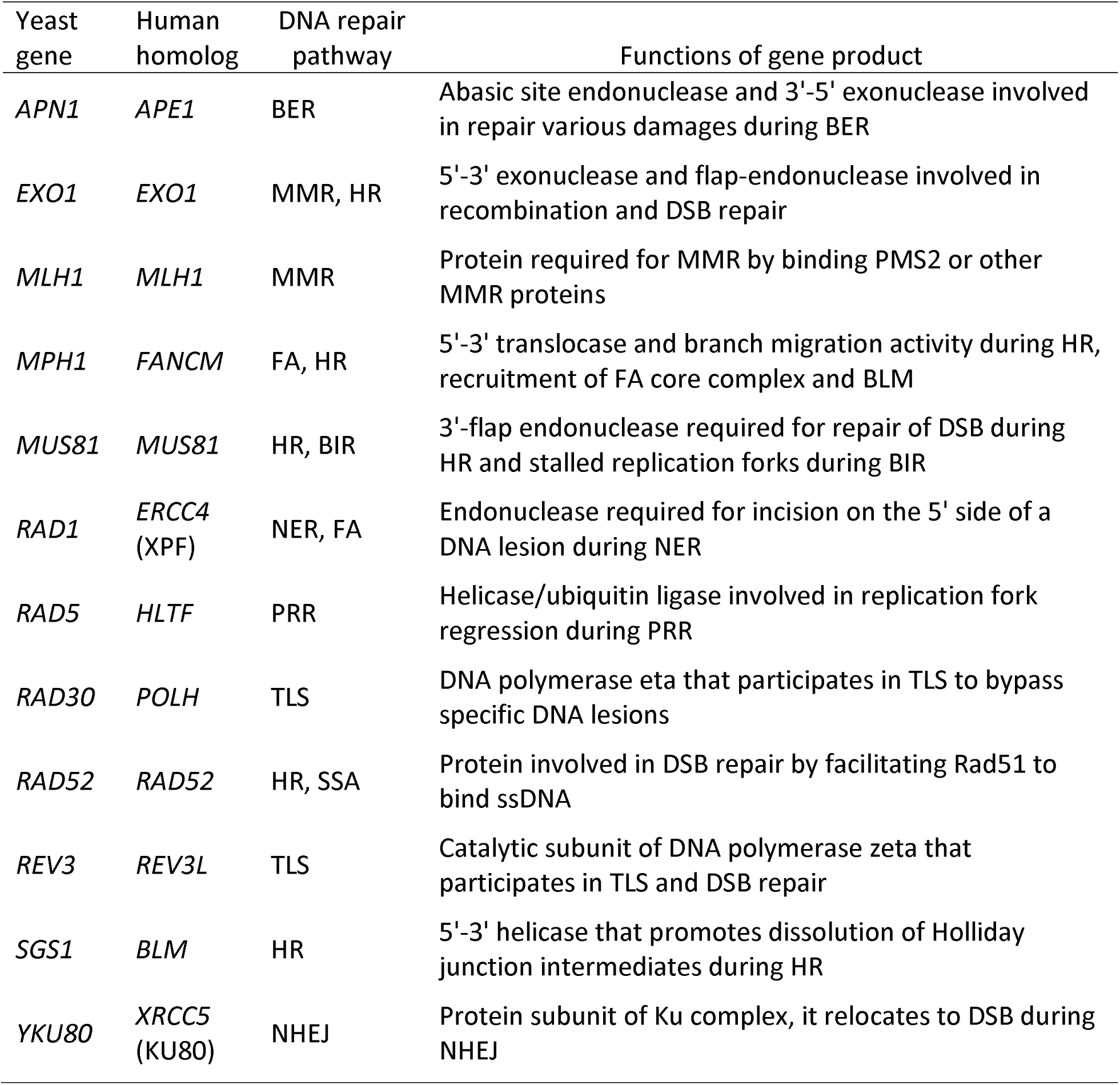
Genes and DNA repair pathways investigated for gene-drug interactions

| Yeast gene | Human homolog | DNA repair pathway | Functions of gene product |
| --- | --- | --- | --- |
| *APN1* | *APE1* | BER | Abasic site endonuclease and 3'-5' exonuclease involved in repair various damages during BER |
| *EXO1* | *EXO1* | MMR, HR | 5'-3' exonuclease and flap-endonuclease involved in recombination and DSB repair |
| *MLH1* | *MLH1* | MMR | Protein required for MMR by binding PMS2 or other MMR proteins |
| *MPH1* | *FANCM* | FA, HR | 5'-3' translocase and branch migration activity during HR, recruitment of FA core complex and BLM |
| *MUS81* | *MUS81* | HR, BIR | 3'-flap endonuclease required for repair of DSB during HR and stalled replication forks during BIR |
| *RAD1* | *ERCC4* (XPF) | NER, FA | Endonuclease required for incision on the 5' side of a DNA lesion during NER |
| *RAD5* | *HLTF* | PRR | Helicase/ubiquitin ligase involved in replication fork regression during PRR |
| *RAD30* | *POLH* | TLS | DNA polymerase eta that participates in TLS to bypass specific DNA lesions |
| *RAD52* | *RAD52* | HR, SSA | Protein involved in DSB repair by facilitating Rad51 to bind ssDNA |
| *REV3* | *REV3L* | TLS | Catalytic subunit of DNA polymerase zeta that participates in TLS and DSB repair |
| *SGS1* | *BLM* | HR | 5'-3' helicase that promotes dissolution of Holliday junction intermediates during HR |
| *YKU80* | *XRCC5* (KU80) | NHEJ | Protein subunit of Ku complex, it relocates to DSB during NHEJ |

Growth rates were measured over a period of 24 hours and the area under the curve was calculated and normalized to the untreated WT to measure fitness (Fig. 1A). Overall, we observed that Cpt and Etp had no significant additional effects on any strain using the concentrations tested and although 5FU inhibited growth of all lines irrespective of the genetic background it only showed a synergistic interaction with *apn1*∆. We also observed that cisplatin had synergistic gene-drug interactions with *rad1*∆, *rad5*∆, *rad52*∆, *rev3*∆ and *sgs1*∆ and that MMS had synergistic interactions with *rad5*∆ and *sgs1*∆ (p<0.05; Fig. 1A). These interactions are consistent with, for example, the known roles for BER in repair of uracil in DNA; or the known complexity of repairing cisplatin interstrand crosslinks (ICLs) which involves at least HR, NER, FA and TLS. Moreover, this screen possibly under-represents chemical sensitivities because very low concentrations of drug were used to allow comparison with fluctuation assays below.

**Figure 1.**
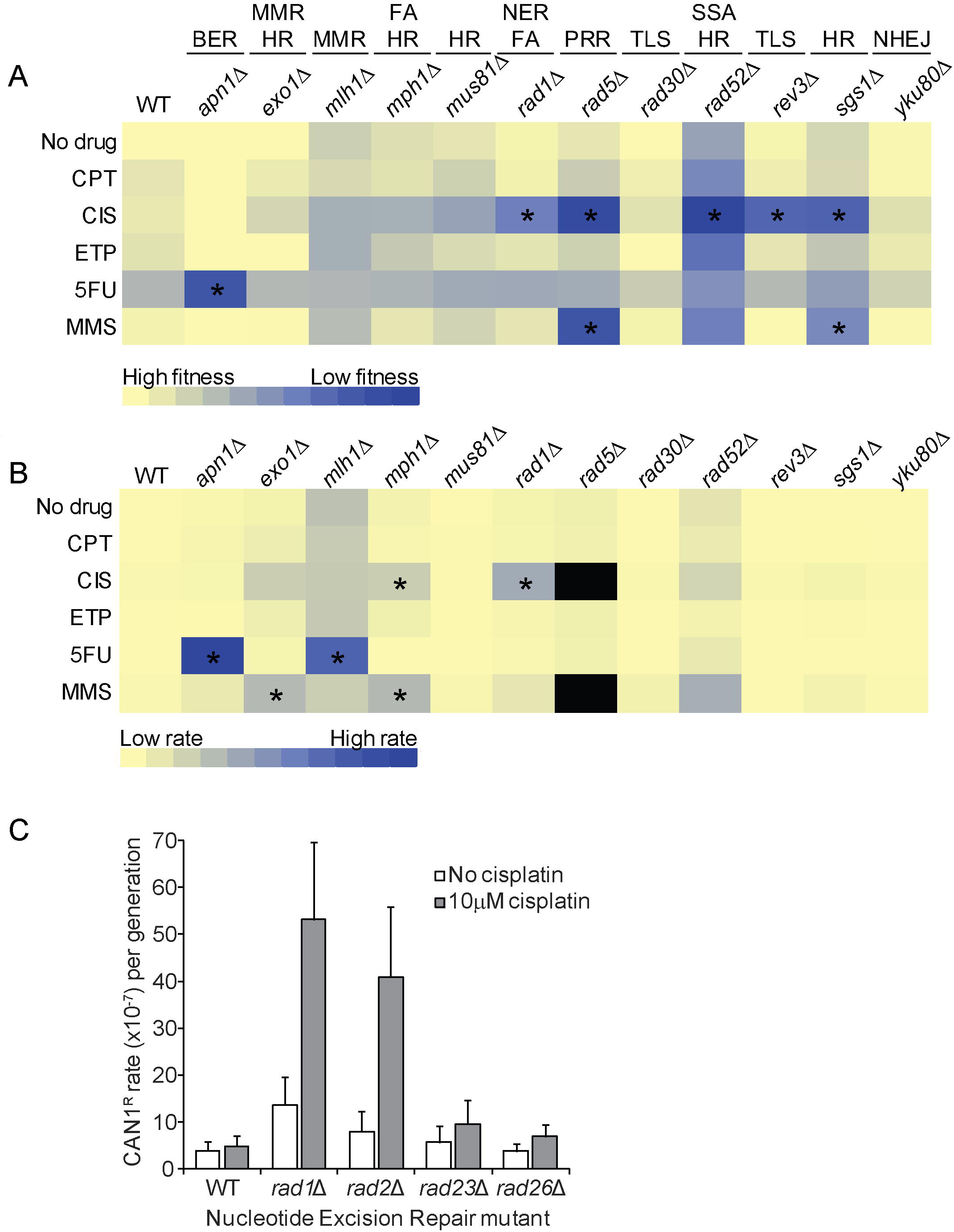
Gene-drug interactions measured by fitness and mutation rates. Heat maps illustrating (A) fitness, and (B) *CAN1* mutation rate, relative to WT. Interactions significantly greater than expected (p>0.05) are indicated *. The transition from yellow to blue indicates greater fitness defects or higher mutation rates. Black boxes for *rad5*Δ indicate not done. (C) *CAN1* mutation rates of other NER deficient strains in cisplatin. *rad1*Δ and *rad2*∆ exhibit SHyp (p<0.05).

Mutation rates of the same matrix were quantified at *CAN1* using a well-plate fluctuation assay (Fig. 1B) (18). In untreated cells the baseline mutation rates matched previously reported rates **(Table S3**). Again we observed that Cpt and Etp had no major mutator effects regardless of the genetic background at the given drug concentrations. On the contrary, cisplatin, 5FU and MMS increased the mutation rates of specific mutants. When this increase in mutation rate exceeded the expected effects of multiplying the effect of the repair deficiency and the effect of the chemical on WT, we defined it as *Synthetic Hypermutation* (SHyp). Applying this metric we identified six cases of SHyp: *mph1*∆-Cis, *rad1*∆-Cis, *apn1*Δ-5FU, *mlh1*∆-5FU, *exo1*∆-MMS and *mph1*∆-MMS (Fig. 1B and **Fig. S1**). Of these cases *rad1*∆-Cis, *apn1*Δ-5FU and *mlh1*∆-5FU showed the most severe SHyp phenotype (17.8, 45.1 and 33.4 relative rates respectively). Importantly, cases of SHyp did not overlap with fitness defects except for *rad1*Δ-Cis and *apn1*Δ-5FU and conversely many strains with fitness defect exhibited no increase in mutation rate. Indeed, survival analysis of cisplatin sensitivity of *sgs1*Δ, *mus81*Δ and *rad1*Δ show nearly identical viability curves over cisplatin concentration but different mutational outcomes (**Fig. S2**). Together our data show that decreased fitness and increased mutability by genotoxins in DNA repair deficient cells are not perfectly linked. This is likely both an issue of dose and DNA repair mechanisms wherein some gene-drug combinations are more likely to elicit toxic or irreparable intermediates.

### The incision step of NER mediates synthetic hypermutation in cisplatin

Rad1 is a nuclease involved in cleaving DNA flanking bulky lesions during NER. Recognition of these lesions can occur during transcription or globally due to the action of upstream NER components that recognize distorted DNA helices (19). To determine which aspects of NER exhibit SHyp with cisplatin we tested mutation rates in three other NER mutants, *rad2*Δ, *rad26*Δ and *rad23*Δ. Deletion of *RAD2* which, like Rad1, encodes a nuclease required for DNA incision and is related to human XPG, also exhibited SHyp with cisplatin (Fig. 1C). However, deletion of *RAD26*, which selectively impairs transcription-coupled (TC) NER, or *RAD23*, which partially impairs global and TC-NER but allows efficient DNA incision, had no SHyp phenotype with cisplatin (Fig. 1C). Since Rad1 acts first, and Rad2 is required in a structural role to position Rad1 (19), loss of either would block incision; consequently, alternative mechanisms of unhooking of the cisplatin damage must be error prone.

### Mutation accumulation and whole-genome sequencing

It is possible that *CAN1* showed variable mutation rates in repair deficient strains due to a locus specific effect. To explore the genomic spectrum of mutations and identify any evidence of such bias, we pulsed diploid WT, *rad1*Δ, *sgs1*Δ and *mus81*Δ with a higher dose of cisplatin and sequenced the whole genomes of 12 independent survivors for each strain (Fig. 2A). Consistent with the *CAN1* fluctuation analysis data, the *rad1*Δ mutant strain acquired significantly more mutations than WT or the other mutants (Fig. 2B). As expected based on the preference of cisplatin for crosslinking guanine residues, the most common mutation types were single-nucleotide variants (SNVs) at C:G basepairs. Moreover, indels seen in *rad1*Δ, which were primarily -1 deletions, occurred at C:G basepairs, and rare dinucleotide substitutions were also evident at motifs containing C:G basepairs (Fig. 2C **and** 2D). To ensure that the rate and type of mutations were due to cisplatin and not simply *rad1*Δ, we passaged diploid WT and *rad1*Δ cells through 40 single-cell bottlenecks and sequenced 12 independent whole genomes after this 1000 generation MA experiment (Fig. 2A). The mutation rate and type evident in the *rad1*Δ MA lines was similar to the *CAN1* rate in untreated *rad1*Δ cells, showed proportionally more mutations at T:A basepairs and no dinucleotide substitutions (**Fig. S3**). No correlations between *rad1*Δ-Cis mutations with transcription, replication timing or leading-strand character were seen, although a weakly significant enrichment (p=0.0103) was seen between nucleosome bound DNA and mutations (20). Thus the specific combination of *RAD1* deletion and cisplatin treatment engages an error prone repair process that causes a global increase in specific types of mutations.

**Figure 2.**
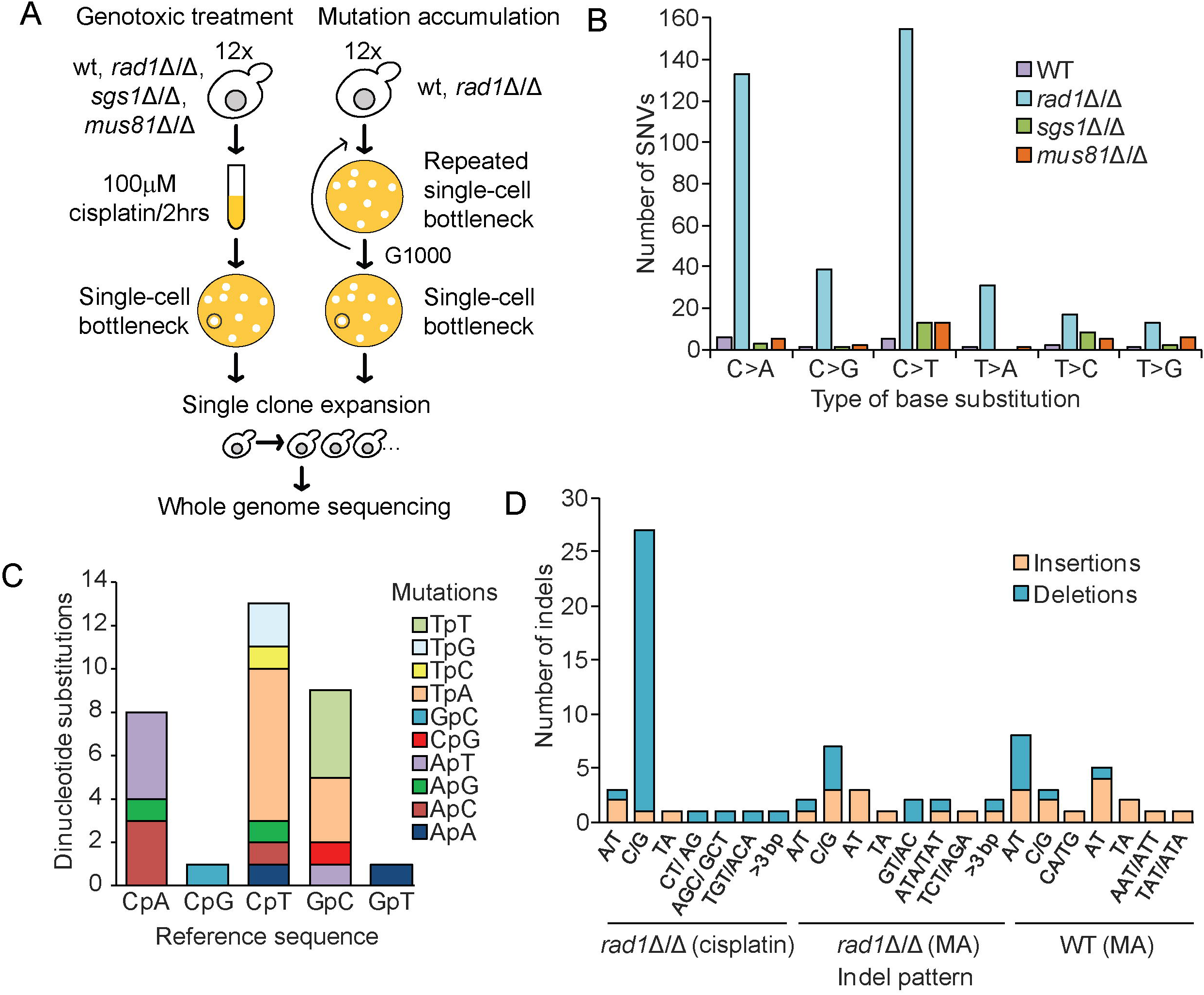
Mutation accumulation and whole genome sequencing. (A) Experimental design of drug treatment and 1000 generation (G1000) approaches for the indicated strains. (B) Summary of WGS results of single nucleotide variants in cisplatin treated WT, *rad1*∆, *sgs1*∆ and *mus81*∆ homozygous diploid strains. SNV number and type are summarized in the bar graphs color-coded as indicated. (C) Dinucleotide substitutions in *rad1*Δ-Cis genomes. The parental dinucleotide sequences are on the x-axis and the mutated sequence is represented by the color of the bar. (D) Pattern of small indels in *rad1*Δ-Cis and MA genomes. Indel size and sequence are on the x-axis and indel types are color coded.

### Cisplatin-rad1Δ mutation signatures suggest 3’ templating of base substitutions

The flanking sequences around mutations can define a signature that is associated with a particular genetic or chemical perturbation. Indeed, previous work in *C. elegans* has examined cisplatin treatment of several mutants including XPF-1 (i.e. *C. elegans RAD1*) deficient animals(11). We extracted flanking sequences for the predominant C>N mutations in cisplatin-treated genomes and performed pLogo and enrichment analysis (Fig. 3A and **Fig. S4A**)(21). This analysis revealed that GpCpT motifs were favored when all C>N mutations were considered. Since cisplatin is expected to preferentially crosslink guanine residues, this signature suggests that ICLs are the mutagenic lesion in yeast *RAD1* mutants. The observed GpCpT motif differs from *C. elegans* where CpCpT motifs were preferentially mutated in *xpf-1* worms (11). Since intrastrand crosslinks are much more common than ICLs (22), our data suggest that *rad1*Δ yeast are either able to repair these in an error-free manner, even when lacking a key NER protein, or that they die and are removed from the experiment. Further pLogo analysis with fixed positions at GC showed that, for each mutation type, adenine was significantly enriched at the -2 position (i.e. ApGpCpT) (Fig. 3A **center and Fig. S4A**). When we analyzed the 3’ position a different picture emerged, the mutation type varied with the +1 base, such that C>T mutations preferentially occurred in GpCpT motifs, and C>A mutations in GpCpA motifs and C>G mutations occurred at GpCpG motifs (Fig. 3B **and Fig. S4B**). Thus, the base which is erroneously inserted at the damaged site is selected by basepairing with the 3’ nucleotide.

**Figure 3.**
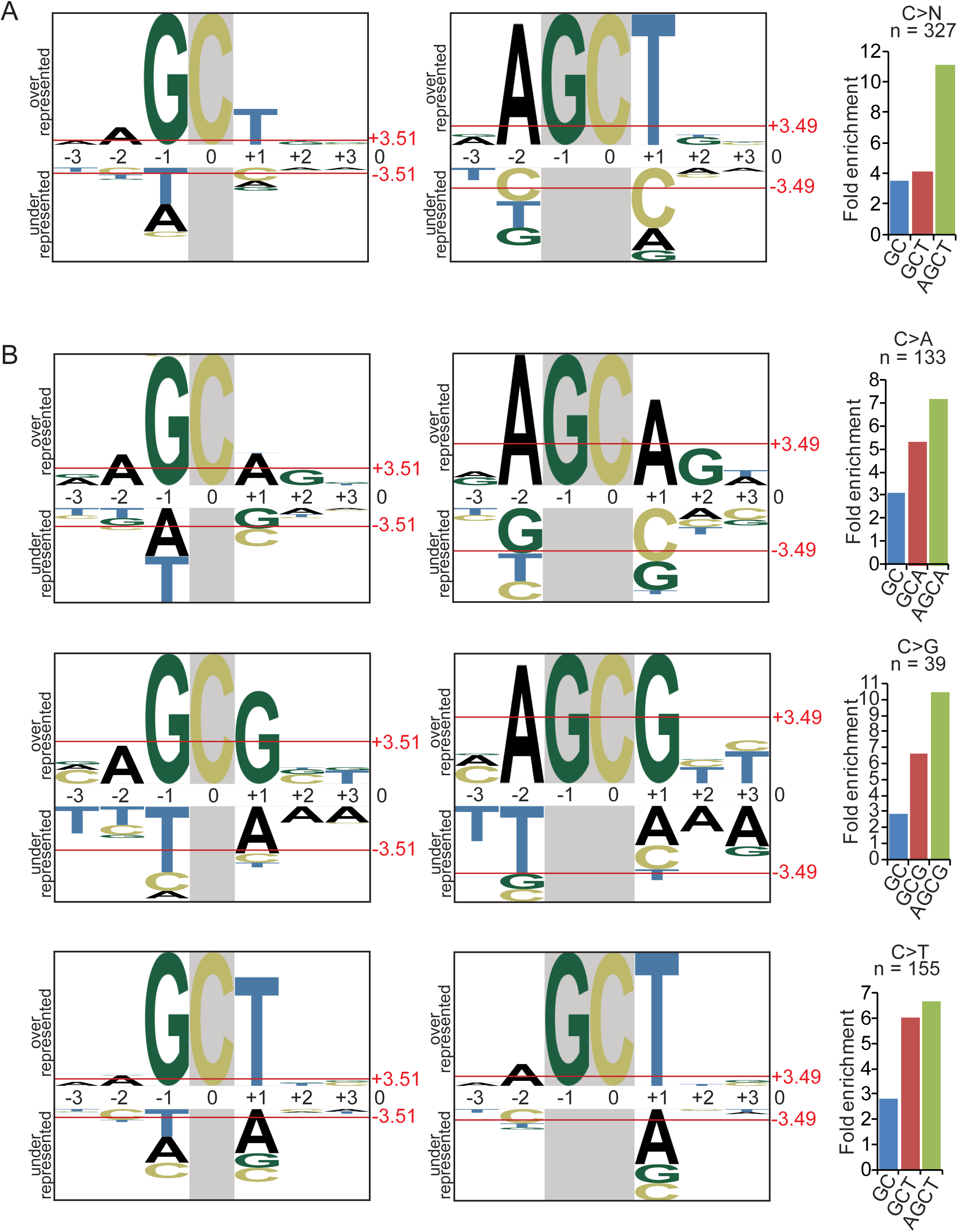
Mutation signature analysis induced by cisplatin in *rad1*∆ diploid yeast. (A) Flanking nucleotides for C>N substitutions are presented as pLogo plots and fold enrichment. The mutated C is centered in each plot with fixed positions at C (left) and GC (center). The red line indicates significant enrichment p<0.05 and the font size indicates the magnitude of enrichment. The increasing fold enrichment for 2- 3- and 4-nucleotide motifs containing a mutated C is shown (left). All analysis was performed with a set of 41-mers and cropped for visualization (see methods) (B) Flanking analysis for C>A, C>G and C>T mutations as presented in A. Note the +1 base changing with the mutation type.

### DNA polymerase ζ drives rad1Δ-cisplatin synthetic hypermutation

Our mutation signature analysis suggests error-prone DNA replication is likely causing mutations in *rad1*Δ cells treated with cisplatin. To identify the relevant polymerase we deleted all enzymatic TLS components individually in the *rad1*Δ background. Loss of *RAD30*, which slightly positively affected fitness when compared with the *rad1*Δ single mutant, showed no effect on *rad1*Δ hypermutation (Fig. 4A and 4B), however additional loss of *REV1* or *REV3* led to inviability when treated with 10μM cisplatin (Fig. 4A). This is consistent with the DNA damage tolerance role of *REV3* and the idea that TLS is supporting viability in *RAD1* deficient cells. A lower dose of cisplatin still revealed SHyp with *rad1*Δ but showed complete suppression of SHyp in *rad1*Δ*rev3*Δ and *rad1*Δ*rev1*Δ cells (Fig. 4B), indicating that Polζ TLS was responsible for the mutations. To determine if endogenous DNA lesions could promote 3’-templated mutations, we scored a published dataset of *URA3* mutations sequenced in unperturbed NER-deficient (*rad14*Δ) cells with or without *TLS* polymerase deletions (23). Remarkably, 3’ templating was observed at 43% of SNVs, deletion of *RAD30* had no effect on this frequency while *REV3* deletion reduced the frequency of SNVs matching the 3’ base to 29% (Fig. 4C). Together these data lead us to propose a model in which slippage and realignment of the DNA template by Rev3 led to the observed mutation signature and is generalizable to endogenous lesions (Fig. 4D and discussed below).

**Figure 4.**
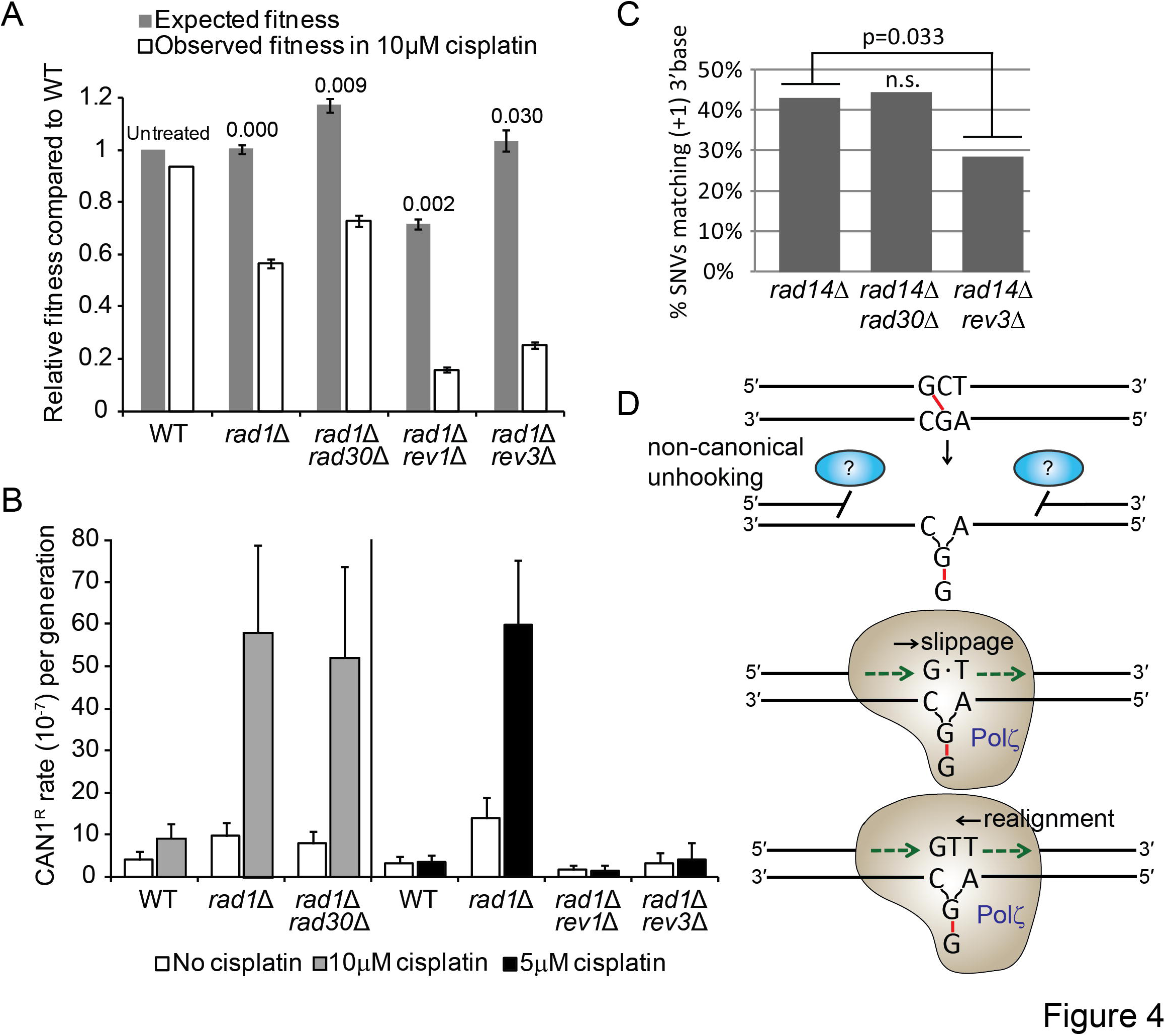
Role of translesion synthesis in hypermutation of cisplatin treated *rad1*∆ cells. (A) Fitness measurements and (B) mutation rates in *rad1*∆ cells lacking the indicated TLS polymerase subunit. (C) Frequency of SNVs identical to 3’ base in *URA3* from the indicated strains from (23). (D) Model for slippage and realignment of DNA polymerase ζ(Rev3) at a cisplatin damaged base. Non-canonical and presumably not optimal unhooking of the ICL creates a substrate for error-prone TLS that promotes slippage of Polζ at the lesion and subsequent realignment upon lesion bypass leading to a 3’-templated SNV.

## DISCUSSION

### Synthetic hypermutation as an alternative measure of phenotypic enhancement in genome maintenance

As fitness datasets on gene-gene and gene-drug interactions accumulate (12, 13), measurements of trigenic interactions (24), or the ways in which chemicals shift genetic-interaction networks (e.g. synthetic cytotoxicity (14, 15)) are enhancing the wiring diagram of the cell. Still other approaches have combined high-content imaging with genetic and chemical perturbations to create conditional phenotypic networks (25). Damage repair and mutation is key to the chemotherapeutic action of alkylating agents and changes in DNA damage repair can have profound effects of therapeutic efficacy. While there is prior evidence that ultra-high mutation rates can drive severe fitness consequences, even lethality (26), the overlap of these phenotypes broadly is not known. By systematically ablating the major DNA repair pathways in yeast and measuring the effects of diverse genotoxic chemicals our data suggest that fitness and chemically induced hypermutation are not necessarily linked.

SHyp interactions between 5FU and *mlh1*Δ and *apn1*Δ are largely consistent with the role of MMR and BER respectively in removing uracil from DNA (27). Indeed, these data show that 5FU can overwhelm the repair capacity of MMR or BER deficient cells without additionally affecting the fitness of MMR impaired cells. This is consistent with evidence that 5FU has multiple modes of action interfering with both DNA and RNA metabolism (27). On the other hand, while both NER and HR are required for repair of cisplatin ICLs and mutants in these pathways had fitness defects, *rad1*Δ but not *sgs1*Δ or other major HR mutants showed SHyp with cisplatin. This suggests that while survival in *rad1*Δ cells is supported by an error prone mechanism, that same pathway is insufficient to support *sgs1*Δ cells which either die, likely due to toxic recombination intermediates, or repair cisplatin damage in an error free way.

### A mutation signature of TLS polymerase slippage and realignment

Whole genome sequencing analysis of mutagenized model organism genomes is a powerful way to link mutation signatures to specific causal phenomena (28). This approach has been used to shed new light on DNA repair mechanism (MMR,(29)), the effects of genotoxic chemicals (11, 30), and the role of cytidine deaminase enzymes in cancer (31). Characterization of our sequencing data revealed that in *rad1*Δ yeast ICLs were the predominant source of mutations since most variants were found at Gp<u>C</u> motifs. This differs from the cisplatin mutation signature of both *C. elegans* and humans where intrastrand crosslinks seem to drive mutagenesis (i.e. mutations at CpC motifs (6, 11)). There are several possible reasons for this including the complexity of the Fanconi Anemia pathway for ICL repair in humans that is only partly represented in yeast, or the high divergence of *REV3* itself (i.e. human REV3L protein is >twice as long, and exhibits only 25% similarity to REV3). While Rev1 plays an important structural role, we predict Rev3 is catalyzing the mutations that we observed in our study, since the catalytic role of Rev1 to insert C opposite the damaged guanine would be error-free and thus not detectable in our assay. Yet another possibility is that intrastrand crosslinks are not repaired in *rad1*Δ cells and lead to cell death, effectively masking the mutations seen at these sites in other systems. We also found evidence of Rev3 slippage and realignment in mutations from untreated cells, suggesting that endogenous, non-cisplatin-induced lesions can promote the same mechanism (23).

Our mutation signature analysis revealed not only a different type of mutagenic lesion in yeast, but also a mechanism involving Polζ, Rev3, which indicates a slippage and realignment event during TLS. Previous work *in vitro* has identified this behavior in another TLS polymerase, human DNA polκ (32). We propose that (Fig. 4D), at least in the specific context of cisplatin adducts, the guanine lesion is bypassed by Rev3 in a manner that uses Watson-Crick basepairing with the undamaged +1 base. This may occasionally lead to an indel, but more often the newly inserted base shifts to mispair with the damaged G residue and is extended with the correct base at the 3’ position. Whether suboptimal DNA incision by compensatory nucleases favor this misincorporation event is an active area of research. Indeed, the nature of DNA incision in the absence of Rad1 is unclear, although structure-specific nucleases such as Slx4, Mus81 and Yen1 have activities that could in principle complement Rad1 loss at some structures (33). Alternatively, new data in *Xenopus* extracts implicates the N-glycosylase activity of NEIL3, an upstream BER factor, in unhooking psoralen and other ICLs (34). Therefore it is also possible that yeast N-glycosylases could act on cisplatin crosslinks in the absence of the preferred Rad1-pathway and create a unique chemical moiety that drives slippage-realignment TLS by Polζ in vivo.

### Synthetic hypermutation and cancer

The framework established here sought to establish the gene-drug relationships driving hypermutation in cells surviving genotoxin treatment beginning with canonical DNA repair pathways and drugs of clinical relevance. There remains a large space to be explored; in yeast there are many hundreds of genes whose loss of function causes genome instability and hundreds of others whose increased dosage causes genome instability (10, 35, 36). The ways in which these will combine with each other and with various chemicals to permit mutagenesis remains incomplete.

Hypermutation of surviving cancer cells after genotoxic chemotherapy could be viewed as negative if it permits the acquisition of chemoresistant mutations. However, current evidence suggests that resistance mutations are often pre-existing in tumor cell populations and their emergence is a clonal selection process rather than chemotherapy induced mutations (37). Indeed, even for therapy-related leukemias, p53 mutations driving leukemogenesis were recently found to be pre-existing when chemotherapy occurred (38). Thus, synthetic hypermutation may not be a major concern for acquiring chemoresistant mutations when treating repair-deficient cancers with genotoxic chemotherapies. While chemotherapy has been shown to minimally impact chemoresistant mutations, the DNA damaging nature of chemotherapeutic agents can increase the overall mutational load (6). We propose that this hypermutation of surviving cells could ultimately be beneficial in contexts where an immune response to tumor neo-antigens is desired as is the case for therapies targeting immune checkpoints (39). The efficacy of immune checkpoint blockade has been correlated with mutational load in lung cancer (40), melanoma (41), and MMR deficient colorectal cancers (42). Understanding more about the interactions of genotoxic chemotherapies with genome maintenance defects in cancer could ultimately support interventions where it is desirable to increase the mutational load or reshape the mutational profile of cancer cells following frontline therapy.

## MATERIALS AND METHODS

### Yeast strains and growth curve analysis

Yeast strains are listed in **Table S1**. Yeast were grown on SD minimal media plus histidine, uracil, leucine, lysine and methionine + 2% dextrose (SD+5) or YPD at 30°C. Growth curves were conducted in triplicate in 96-well plates using a TECAN M200 plate reader at 30°C, with OD600 measured every 30 minutes (35). The area under the growth curve was used as a measure of fitness and compared to the expected fitness value (multiplicative model) using a t-test(35). Unless otherwise indicated, drug concentrations were determined empirically by permitting well saturation of *rad52*Δ cells over 3 days, to be 1 μM Cpt, 10 μM cisplatin, 200 μM Etp, 10 μM 5FU and 0.0005% MMS.

### Fluctuation analysis and synthetic hypermutation analysis

Fluctuation analysis at *CAN1* was performed in 96-well plates, with a minimum of 18 replicates per condition, exactly as described(18), except that cells were grown to saturation in the presence of drugs in SD+5 for 48 hours at 30°C. Average cell numbers in each well were determined using a TC-20 cell counter (BioRad), and canavanine-resistant colony counts were converted to mutation rates using the Ma-Sandri-Sarkar maximum-likelihood estimator calculator (43). Synthetic hypermutation was calculated as:

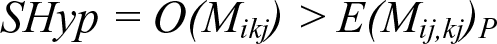

Where *M_i_* is the mutation rate of mutant *i; M_k_* is the mutation rate caused by drug *k* in WT; *M_ik_* is the mutation rate caused by drug *k* in mutant *i*; each are expressed as *_j,_* the fold increase over untreated WT, to give observed mutation rates *O(M)*. *E(M_ij,kj_)_P_* is the expected mutation rate as the product model that is defined as *E(M_ij,kj_)_P_ = M_ij_ · M_kj_*. Alternatively, the additive expected mutation rate was also considered and is defined as *E(M_ij,kj_)_A_ = M_ij_ + M_kj_* (**Fig. S1**). Synthetic hypermutation was considered if the lower 95% confidence interval of the observed mutation rate fell above the expected value.

### Mutation accumulation and whole-genome sequencing

Diploid strains were pulsed with 100 μM cisplatin for 2 hours before plating survivors on YPD agar. For MA lines single colonies were plated on YPD and passaged 40 times by restreaking to fresh YPD every 2-3 days (~1000 generations). 12 independent lines for cisplatin treated or MA were outgrown for genomic DNA extraction (10). DNA was subjected to 125 bp paired end whole genome sequencing at Canada’s Michael Smith Genome Sciences Centre using the Illumina HiSeq2000 platform. The average sequencing coverage compared to the haploid reference was 129X (i.e. >60x for each diploid genome). Sequence files are deposited at the Sequence Read Archive (http://www.ncbi.nlm.nih.gov/sra) (SRP091984). Reads were aligned to the Saccer3 reference genome using Burrow-Wheeler Aligner 0.5.7 (44). Variants were identified exactly as described previously (10). Mutations were extracted by comparing evolved to parental genomes. Variant calling was carried out using the mpileup utility of the SAMtools package 0.1.18 (45) with a threshold of 20. Copy-number variants were analyzed using a locally modified version of CNAseq (46). Variant calls were manually confirmed with the Integrated Genomics Viewer (Broad Institute).

### Mutation signature analysis

Flanking sequence was extracted from the Saccer3 genome based on the mutation positions (**Table S2**). The enrichment of motifs within the diploid *rad1*Δ+cisplatin genomes was calculated as described (21). Briefly, enrichment is the ratio of the number of mutations at a motif multiplied by the total number of C or G bases in 41mers (+/- 20bp) flanking the mutation, divided by the number of mutations of that type (e.g. C>N) multiplied by the total number of motifs present in the 41mers. The significance of these enrichments was calculated using the hypergeometric distribution. pLogos were generated online http://plogo.uconn.edu/ (47). As foreground input n(fg), 41-mer sequences from the *rad1*∆-cisplatin genomes were used. These sequences were centered at the mutated C for all substitutions with fixed positions at <u>C</u> and G<u>C</u>. As background input n(bg), unique 41-mer sequences automatically generated by the pLogo tool from the Saccer3 yeast genome were used. Reanalysis of variants from Stone et al. was performed by manually scoring whether the SNV type matched the base immediately 3’ of the mutation (23). The proportions of these 3’-templated SNVs were compared with a Fisher exact test.

## ACKNOWLEDGEMENTS

We thank Nigel O’Neil and members of the Stirling lab for reading the manuscript and Philip Hieter for yeast strains. P.C.S. is a Canadian Institutes of Health Research (CIHR) New Investigator and a Michael Smith Foundation for Health Research Scholar. The work is funded by CIHR (MOP-136982) and Natural Sciences and Engineering Council of Canada (RGPIN 2014-04490) grants to P.C.S. and funds from the BC Cancer Foundation.

